# White Matter Correlates of Impulsive Behavior in Healthy Individuals: A Diffusion MRI Study

**DOI:** 10.1101/2023.10.14.562328

**Authors:** Fatemeh Rashidi, Mohammadamin Parsaei, Iman Kiani, Arash Sadri, Mohammad Hadi Aarabi, Seyed Reza Darijani, Yune Sang Lee, Hossein Sanjari Moghaddam

## Abstract

**Background:** Impulsivity is closely related to the tendency to engage in risky behaviors. Previous research identified macrostructural brain alterations in individuals exhibiting impulsive behaviors. Understanding the microstructural brain changes linked to impulsivity can elucidate its underlying mechanisms and guide effective treatment strategies. In this study, we employed diffusion magnetic resonance imaging (DMRI) connectometry to investigate white matter tracts associated with impulsivity while considering potential sex differences.

**Methods:** We enrolled 218 healthy participants from the Leipzig Study for Mind-Body-Emotion Interactions (LEMON) database. Correlations between DMRI-derived white matter changes and impulsivity were assessed using scores from the UPPS Impulsive Behavior Scale’s four subscales (lack of perseverance (PE), lack of premeditation (PM), sensation seeking (SS), and negative urgency (NU)).

**Results:** Our findings revealed negative correlations between quantitative anisotropy (QA) values in bilateral cerebellum, middle cerebellar peduncle (MCP), and the severity of PE and PM across the cohort. Additionally, QA values within MCP, corpus callosum (CC) body, and forceps major exhibited negative correlations with SS. Conversely, QA values in forceps minor were positively correlated with PM, and QA values in both the forceps minor and bilateral cingulum showed positive correlations with SS. Remarkably, the observed correlations between UPPS subscale scores and QA value alterations within white matter tracts varied between males and females.

**Conclusions:** Impulsivity is correlated with discernible alterations in white matter integrity across diverse tracts, including CC, cerebellum, and cingulum. Moreover, males and females show distinct patterns of correlations between white matter integrity and impulsivity.

**Highlights:**

- Impulsivity is associated with QA changes and white matter alterations in various brain tracts.
- Lower white matter integrity in the cerebellum is negatively correlated with impulsivity.
- QA values in the CC parts showed divergent correlations with impulsive behavior.
- The patterns of correlations across various tracts varied between males and females.

## 1. Introduction

Impulsivity is a complex trait that is typically defined as the tendency to engage in immediate actions without adequate consideration of potential outcomes or consequences (Dalley and Robbins, 2017). It is characterized by an array of deficits in areas such as delayed gratification, impulse and urge control, decision-making, and maladaptive behaviors (Robbins, Curran and de Wit, 2012). Concordantly, it is associated with heightened propensities for risky decision-making, substance use disorders, eating disorders, and suicide attempts (Barati et al., 2020; Dawe and Loxton, 2004; Parsaei et al., 2023), signifying an enhanced vulnerability and diminished quality of life among individuals exhibiting impulsive behaviors. Noteworthy, impulsivity has been marked by linguistic and conceptual ambiguities, making its characterization more challenging.

Self-reported questionnaires, such as the Urgency-Premeditation-Perseverance-Sensation (UPPS) Impulsive Behavior Scale, are commonly employed to assess impulsivity (Whiteside and Lynam, 2001). This scale is based on the five-factor model of personality and consists of four subscales: negative urgency (NU), lack of premeditation (PM), lack of perseverance (PE), and sensation seeking (SS) (McCrae and John, 1992). The UPPS Scale was developed by conducting an exploratory factor analysis on various impulsivity measures which resulted in identifying these four distinct yet interconnected personality facets of impulsivity (Whiteside and Lynam, 2001). Rather than mere variations of impulsivity, these facets are understood as distinct psychological processes that can incline individuals to engage in behavior without fully considering their possible negative outcomes (Whiteside and Lynam, 2001). The NU refers to the tendency to show impulsive behaviors in response to negative emotions. The PM denotes the tendency to behave without considering the potential consequences of the behavior. The PE captures the difficulty of staying focused on a task, particularly if it is tedious or challenging, and SS refers to the tendency to seek out novel and thrilling experiences(Whiteside and Lynam, 2001). Prior research has demonstrated that different types of impulsive behavior are linked to distinct and specific impulsivity traits (Kale, Stautz and Cooper, 2018; Masoudi et al., 2021; Rømer Thomsen et al., 2018). For instance, Bousardt et al. (2018) observed that NU is associated with aggressive behaviors, while PM is linked to SS, and PE is associated with problematic alcohol use (Bousardt et al., 2018).

There also has been considerable research on the neural basis of impulsive behavior. Magnetic resonance imaging (MRI) sequences have revealed a discernible link between impulsivity and structural alterations in grey matter volume across diverse brain regions, encompassing the frontal, parietal, temporal, and occipital cortices (Chen et al., 2022b). Moreover, diffusion tensor imaging (DTI) sequences have uncovered alterations in white matter microstructure among individuals exhibiting impulsivity traits (Ikuta, Del Arco and Karlsgodt, 2018; Moeller et al., 2005; Rashidi et al., 2023). However, the observations on exactly how the white matter microstructure alters in relation to impulsivity have been seemingly inconsistent and, therefore, controversies have ensued. Some studies have reported reduced white matter integrity, as measured by fractional anisotropy (FA), within some white matter tracts (Moeller et al., 2005), while others have reported increased integrity within some other white matter tracts (Ikuta, Del Arco and Karlsgodt, 2018). Collectively, there seems to be a substantive alteration in the white matter tracts of impulsive individuals; however, the direction in which each tract gets altered is still unknown.

Diffusion MRI (DMRI) is an MRI method that enables the investigation of the brain microstructure by analyzing the diffusion of water within its tissues (Chilla et al., 2015). Connectometry is a novel approach to analyzing DMRI data in which the water diffusion density is quantified within a voxel in various directions. This approach aims to find similarities in local connectivity patterns and map white matter tracts (Yeh, Badre and Verstynen, 2016). Unlike traditional DTI, connectometry emphasizes water diffusion density over diffusivity or speed in specific directions (Yeh, Badre and Verstynen, 2016). This emphasis enhances spatial resolution, enabling the identification of white matter fibers even in regions with intricate crossing or intersecting tracts, such as long associational white matter fibers (Huisman, 2010). Utilizing a predefined atlas of diffusion density, the spin distribution function (SDF) is computed for each white matter voxel in various directions (Yeh, Badre and Verstynen, 2016). SDF can be converted into quantitative anisotropy (QA), a density-based index, which can be used to extract fiber tracts, determine between-group differences, and establish correlations between diffusion density and variables of interest (Sobhani et al., 2019; Yeh, Tang and Tseng, 2013).

The present study investigates the relationship between the microstructural integrity of white matter fiber tracts and impulsivity, as assessed by four subscales of UPPS scores, in healthy individuals. Moreover, taking into account previous research findings that have highlighted distinctions in impulsive behavior between males and females (Dretsch and Tipples, 2011) and sex differences in brain structures among individuals exhibiting impulsivity (Kogachi et al., 2017), we performed additional analyses to examine the relationships between white matter connectometric measures and UPPS scores separately for males and females.

## 2. Methods

### 2.1. Study data

The data used in this research were obtained from the Leipzig Study for Mind-Body-Emotion Interactions (LEMON) dataset (Babayan et al., 2019). The LEMON dataset comprises two groups of young (20-35 years old) and old (59-77 years old) adults. The primary purpose of LEMON was to investigate the intricate interplay between mind, body, and emotions (Babayan et al., 2019). This dataset provides a comprehensive collection of information, encompassing psychological evaluations, emotional and personality assessments (including the UPPS inventory), psychiatric evaluations, and physiological measurements (including DMRI data) from 227 healthy participants, collected from September 2013 until September 2015 in Leipzig, Germany (Babayan et al., 2019).

In the development of the LEMON dataset, individuals who fulfilled any of the following criteria were excluded: (1) diagnosis of hypertension without intake of antihypertensive medication, (2) diagnosis of any other cardiovascular disease, (3) history of any psychiatric disorder that required inpatient treatment for longer than two weeks, within the last ten years, (4) history of neurological disorders, (5) history of malignant diseases, (6) intake of any of the following medications: centrally active medications, beta- and alpha-blockers, cortisol, any chemotherapeutic or psychopharmacological medications, (7) positive drug anamnesis (extensive alcohol, methyl enedioxymethamphetamine, amphetamine, cocaine, opiates, benzodiazepine, and cannabis), (8) MRI exclusion criteria (metallic implants, braces, non-removable piercings, tattoos, pregnancy, claustrophobia, tinnitus, and any surgical operation in the last 3 months), (9) previous participation in any scientific study within the last 10 years, and (10) previous or current enrollment in undergraduate, graduate or postgraduate studies.

### 2.2. Participants

218 participants of the LEMON dataset were included in this study and their DMRI data and UPPS scores were extracted. The original study conducted for compiling the LEMON dataset was carried out according to the Declaration of Helsinki (Association, 2013) and was approved by the Ethics Committee of the University of Leipzig (reference number 154/13-ff).

### 2.3. UPPS Impulsive Behavior Scale

UPPS was developed with the objective of assessing impulsivity, achieved through the computation of four distinct measures, namely: (1) negative urgency (NU), which captures the propensity to engage in hasty actions as a response to intense negative emotions; (2) lack of premeditation (PM), entailing impulsive behaviors executed without prior consideration; (3) lack of perseverance (PE), indicative of a disposition to leave tasks incomplete; and (4) sensation seeking (SS), encompassing proclivities towards pursuing novel and stimulating experiences (Whiteside and Lynam, 2001). Each individual item within the assessment is evaluated using a four-point Likert scale, ranging from 1 (strongly agree) to 4 (strongly disagree), thereby indicating the participant’s concurrence with the provided statements. Notably, every subscale consists of a range of 10 to 14 items, and a higher aggregate score within each subscale signifies an elevated degree of impulsivity.

### 2.4 Diffusion MRI acquisition

Axial whole-brain high angular resolution DMRI images were acquired using a 3-Tesla Siemens MAGNETOM Verio device (Siemens Healthcare GmbH, Erlangen, Germany) equipped with a 32-channel head coil. The images were acquired with 60 diffusion directions, 1000 s/mm^2^ b-value, 80 ms echo time, 7000 ms repetition time, 1502 Hz/pixel bandwidth, 0.78 ms echo spacing, 220 mm field of view, 1.7 mm isotropic voxel dimension, and 128×128 matrix. Additional details regarding the DMRI acquisition protocol can be found in the article by Babayan et al. (Babayan et al., 2019).

### 2.5. Diffusion MRI processing and DMRI connectometry

The DMRI data were corrected for subject motion, eddy-current distortions, and susceptibility artifacts due to the magnetic field inhomogeneity using the ExploreDTI toolbox (Leemans et al., 2009). DMRI connectometry analyses were performed using the software DSI Studio (Online address: https://dsi-studio.labsolver.org). Using the q-space diffeomorphic reconstruction (QSDR), DMRI data were reconstructed in the Montreal Neurological Institute (MNI) space to obtain the SDF value which is the peak distribution value for each voxel orientation (Yeh and Tseng, 2011). The SDF is subsequently transformed into QA, which is utilized to create a connectivity matrix of all the voxels within the region of interest (Yeh et al., 2013). The QSDR algorithm is a model-free approach that generates a matrix of orientation functions at varying diffusing spins, which allows for the quantification of the density of water diffusion in different directions for each voxel (Razek et al., 2018). This provides DMRI connectometry with greater spatial resolution and statistical power for fiber tracking than conventional DTI methods (Razek et al., 2018).

Local connectomics, as delineated by Yeh et al. (2018), involves the utilization of QA computations to determine water diffusion density for a given fiber orientation (Yeh et al., 2018). This methodology subsequently facilitates the examination of white matter tracts, enabling comparative analysis among different groups and the identification of associations between white matter fibers and various variables. The present study employed the DSI Studio software in conjunction with the DMRI connectometry protocol to investigate associations between the QA values of white matter tracts and the severity of impulsivity. Local connectomes were selected using a deterministic fiber tracking algorithm that followed the core pathway of each fiber bundle.

### 2.6. Statistical analysis

To assess the association between impulsivity and calculated QA measures, a regression analysis employed. First, the regression analysis encompassed the entire sample, investigating the interplay between QA values of white matter tracts and four distinct UPPS subscales while controlling for age, sex, and overall UPPS scores of the participants as covariates. Subsequently, the analysis was separately conducted on male and females, with their age and total UPPS scores being controlled. Subsequently, based on the recommendations from the software developer, obtained p-values were subject to correction for multiple comparisons using the false discovery rate (FDR) method. The computation of the FDR involved the application of 2000 random permutations to the group labels. This process served to establish the null distribution of tract lengths, enabling the derivation of an accurate FDR estimate. The employment of permutation tests facilitated the control and correction of type-1 error inflation arising from multiple comparisons. Finally, an FDR threshold of less than 0.05 was adopted for tract selection. This criterion guided the reporting of tracts exhibiting statistically significant correlations. This criterion guided the reporting of tracts exhibiting statistically significant correlations.

## 3. Results

### 3.1. Participant characteristics

A total of 218 participants with a mean age of 39.15 years (SD = 20.18) and a male-to-female ratio of 140/78 were enrolled. Table 1 illustrates the demographic characteristics of the sample and the mean scores of four UPPS subscales in the whole sample, males, and females.

**Table 1.** Demographic characteristics of the participants.

|  | All participants<br>(n=218) | Male (n=140) | Female (n=78) | P-value |
| --- | --- | --- | --- | --- |
| Age | 39.15 (20.18) | 36.17 (18.81) | 44.48 (21.56) | 0.005 |
| Handedness |  |  |  | 0.014 |
| Right | 193 (88.5%) | 124 (88.6%) | 69 (88.5%) |  |
| Left | 21 (9.6%) | 16 (11.4%) | 5 (6.4%) |  |
| Ambidextrous | 4 (1.8%) | 0 (0%) | 4 (5.1%) |  |
| BMI | 24.24 (3.65) | 24.38 (3.47) | 23.99 (3.98) | 0.460 |
| Education |  |  |  | 0.604 |
| None | 2 (0.9%) | 1 (0.7%) | 1 (1.3%) |  |
| Gymnasium | 169 (77.5%) | 105 (75%) | 64 (82.1%) |  |
| Hauptschule | 7 (3.2%) | 5 (3.6%) | 2 (2.6%) |  |
| Realschule | 40 (18.3%) | 29 (20.7%) | 11 (14.1%) |  |
| Smoking |  |  |  | 0.349 |
| Non-smoker | 181 (83%) | 117 (83.6%) | 64 (82.1%) |  |
| Occasional smoker | 21 (9.6%) | 11 (7.9%) | 10 (12.8%) |  |
| Smoker | 16 (7.3%) | 12 (8.6%) | 4 (5.1%) |  |
| UPPS-Lack of Perseverance | 25.74 (4.85) | 25.41 (4.75) | 26.34 (5.00) | 0.175 |
| UPPS-Lack of Premeditation | 22.52 (3.92) | 22.58 (3.95) | 22.42 (3.90) | 0.770 |
| UPPS-Sensation Seeking | 19.18 (4.67) | 19.92 (4.90) | 17.85 (3.91) | 0.002 |
| UPPS-Urgency | 32.41 (7.02) | 34.30 (6.59) | 29.02 (6.51) | <0.001 |

### 3.2. Correlations between UPPS and DMRI connectometry

Table 2 presents an overview of the correlations observed between the QA values of the white matter tracts and the UPPS subscales. In the entire sample, we observed a negative correlation between the QA values of the middle cerebellar peduncles (MCP) and the bilateral cerebellum and the deficiencies in PE and PM (FDR = 0.0004 for both). Furthermore, the QA values within the MCP, as well as the body and forceps major of the corpus callosum (CC) display negative correlations with SS (FDR = 0.0001). Conversely, the QA values associated with the white matter in the forceps minor exhibit a positive correlation with PM (FDR = 0.004), while the QA values within both the forceps minor and the bilateral cingulum manifest positive correlations with SS (FDR = 0.0005).

**Table 2.** Correlations between UPPS subscales and the QA values in white matter tracts.

| Analysis group | UPPS subscale | Positive correlations | Negative correlations |
| --- | --- | --- | --- |
| Entire cohort | Lack of Perseverance | NS | MCP and bilateral cerebellum (FDR=0.0004) |
|  | Lack of Premeditation | Forceps minor (FDR=0.004) | MCP and bilateral cerebellum (FDR=0.0004) |
|  | Sensation Seeking | Forceps minor and bilateral cingulum (FDR=0.0005) | MCP, body of CC, and forceps major (FDR=0.0001) |
|  | Urgency | NS | NS |
| Male cohort | Lack of Perseverance | NS | Forceps minor and MCP (FDR=0.001) |
|  | Lack of Premeditation | Forceps minor and major (FDR=0.002) | MCP and L. cerebellum (FDR=0.0001) |
|  | Sensation Seeking | Tapetum of CC (FDR=0.005) | MCP and fornix (FDR=0.003) |
|  | Urgency | Bilateral cingulum (FDR=0.0005) | Forceps major, tapetum of CC, R. cerebellum (FDR=0.001) |
| Female cohort | Lack of Perseverance | R. medial lemniscus, R. CST, tapetum of CC (FDR=0.013) | L. cerebellum and MCP (FDR=0.0004) |
|  | Lack of Premeditation | Body and splenium of CC, and bilateral cingulum (FDR=0.004) | L. cerebellum (FDR=0.005) |
|  | Sensation Seeking | NS | NS |
|  | Urgency | NS | Forceps minor, R. ATR, L. cingulum (FDR=0.005) |

**ATR:** anterior thalamic radiation, **CC:** corpus callosum, **CST:** corticospinal tract, **FDR:** false discovery rate, **L:** left, **MCP:** middle cerebellar peduncle, **NS:** non-significant, **R:** right, **QA:** quantitative anisotropy

In the male cohort, we observed statistically significant negative correlations between these: PE and the QA values within the forceps minor and the MCP (FDR = 0.001); PM and the QA values in the MCP and the left cerebellum (FDR = 0.0001); SS and the QA values of the MCP and the fornix (FDR = 0.003); and NU and the QA values in the forceps major, the tapetum of CC, and the right cerebellum (FDR = 0.001). On the other hand, we observed positive correlation between PM and the QA values in the forceps minor and major (FDR = 0.002), and also, between SS and NU and the QA values in the tapetum of CC and the cingulum, respectively (FDR = 0.005 for both).

In the female cohort, we observed statistically significant negative correlations between these: PE and the QA values of the left cerebellum and the MCP (FDR=0.0004); PM and the QA values of the left cerebellum (FDR=0.005); and NU and the QA values within the forceps minor, the right anterior thalamic radiation (ATR), and the left cingulum (FDR=0.005). On the other hand, we observed a positive correlation between the QA value of the right medial lemniscus, the right corticospinal tract (CST), and the tapetum of the CC and PE (FDR = 0.013). Similarly, there was a positive correlation between PE and the QA values of the body and the splenium of the CC, as well as the bilateral cingulum (FDR=0.004) (Fig. 1).

**Figure 1.**
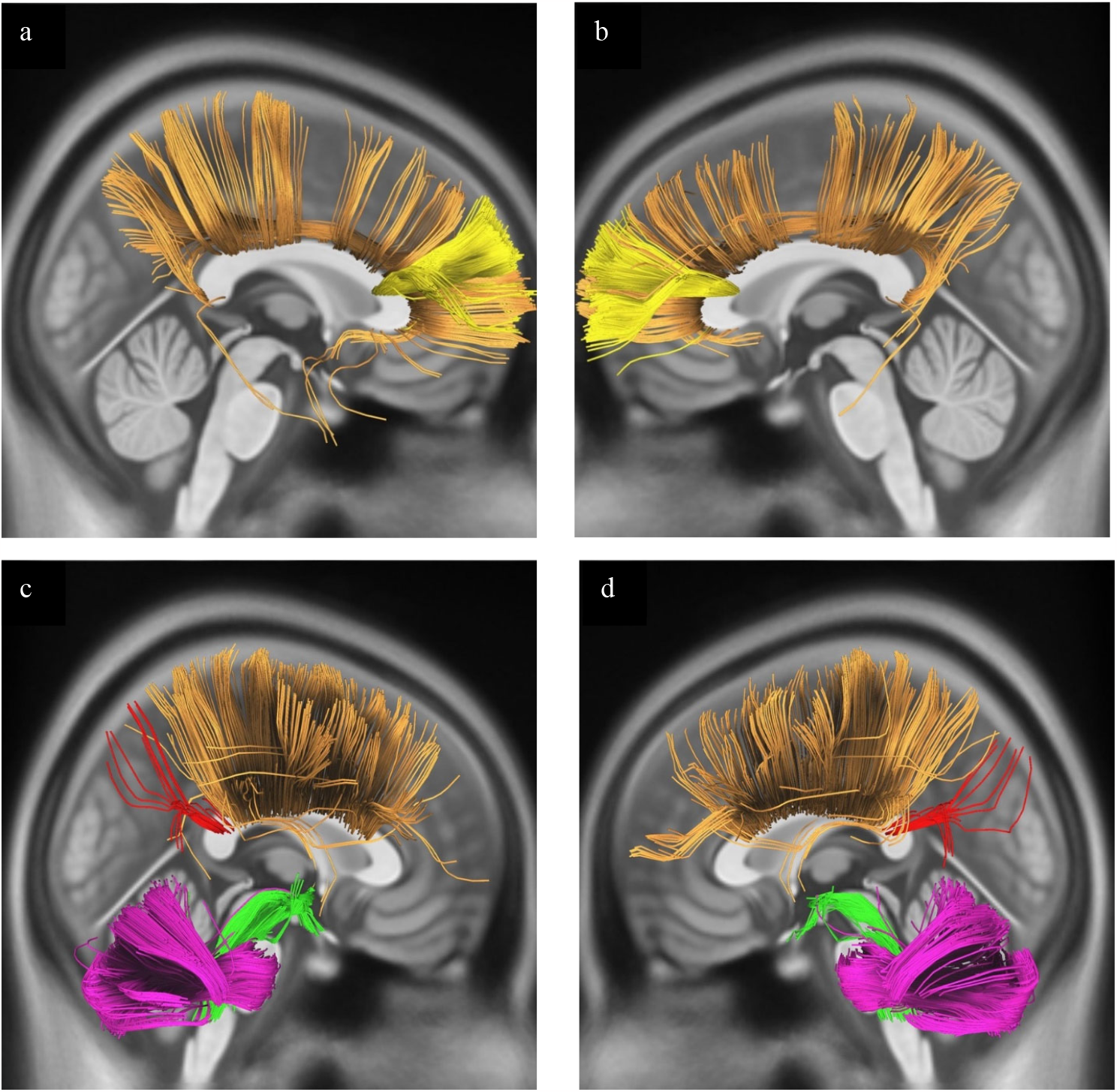
White matter tracts associated with UPPS subscales in entire cohort. a and b) Bilateral cingulum (orange) and forceps minor (yellow). c and d) MCP (green), bilateral cerebellum (pink), body of CC (orange), and forceps major (red).

## 4. Discussion

In the present study, we investigated the association between the microstructural integrity of brain circuits and impulsive behavior, evaluated by the UPPS subscales, in a sample of healthy adults. Based on our findings, PE and PM appears to be negatively correlated with the integrity of the MCP and the bilateral cerebellum. Also, PE showed a positive association with the QA values in the forceps minor. Furthermore, SS seems to be positively correlated with the QA values in the forceps minor and the bilateral cingulum, and negatively correlated with the QA values in the MCP, the body of CC, and the forceps minor. Additionally, our subgroup analysis revealed that males and females had distinct correlation patterns between their white matter tract integrities and the UPPS scores. To our knowledge, this is the first study using DMRI connectometry to explore the brain white matter alterations associated with impulsive behaviors.

Our most consistent observation was the negative correlation between the QA values in the cerebellar regions (cerebellum and MCP) and the degree of impulsiveness in the entire cohort, as well as the males and the females. This is in line with previous research that has reported decreased white matter integrity within the anterior lobe of the left cerebellum in schizophrenic patients (Wei et al., 2011). Also, patients with cerebellar ataxia demonstrate a higher prevalence of impulsive behaviors (Chen et al., 2022a; Lai et al., 2023). Moreover, altered frontocerebellar functional connectivity has been observed in patients with higher impulsivity (Jung et al., 2014; Piccoli et al., 2020). These findings are in line with the observations and hypothesis that the cerebellum has a modulatory role in preventing immediate and unplanned behaviors by inhibiting the prefrontal cortex (Miquel et al., 2019). Based on this hypothesis, the cerebellum regulates ongoing behaviors in the face of changing environmental conditions by modifying the prefrontal reaction to the incoming external and internal stimuli (Miquel et al., 2019). Overall, our findings supported earlier research on the critical role of cerebellar regions in maintaining balance in individuals’ behavioral activities, which explains its lower integrity in persons with higher impulsive tendencies.

Our findings reveal a notable link between white matter integrity within the CC tract and impulsivity. Within the entire cohort, the degree of sensation seeking has a negative correlation with the QA values in the body and the forceps major of the CC. This concurs with the prior investigations that have highlighted an inverse relationship between impulsivity and white matter integrity, as measured by FA, in the genu and the body of the CC (Liu et al., 2010; Sadek et al., 2021). CC is a neural bridge that connects the cerebral hemispheres and facilitates the transmission of visual, auditory, and somatosensory information to the regions accountable for higher-order cognitive functions (Goldstein et al., 2023). Comprising four components – forceps major (splenium), body, genu, and forceps minor (rostrum) – the CC plays a pivotal role in interhemispheric communication (Goldstein et al., 2023). The fibers of the genu traverse and give rise to the forceps minor and connect the frontal cortices. Those of the splenium extend posteriorly to form the forceps major, linking the occipital lobes (Goldstein et al., 2023). Transversely traversing the cerebral cortex, the body fibers give rise to the corona radiata and other substantial white matter pathways (Goldstein et al., 2023). Lastly, rostral fibers connect the orbital regions of the frontal lobes (Goldstein et al., 2023). Given the vital role of CC in transferring emotion-related data, it is reasonable to expect that individuals exhibiting impulsive behaviors might manifest altered CC structures (Lischke et al., 2017).

The existing literature has predominantly indicated that the white matter integrity decreases within the CC of individuals showing impulsive behaviors; however, conflicting observations have also been reported. For instance, Treit et al. (2014), Alfano et al. (2021), and Stansberry et al. (2022) have reported that individuals with higher FA values in the corpus callosum tend to exhibit greater impulsivity (Alfano et al., 2021; Stansberry, Willliams and Ikuta, 2022; Treit et al., 2014). These observations are in line with our results which have revealed a positive correlation between the integrity of the forceps minor and the degree of sSS and PM in the entire cohort. However, our subgroup analysis led to divergent findings regarding the correlations between the integrity of the CC and impulsivity in males and females. In the male cohort, the QA values in the tapetum of the CC had positive correlations with SS and negative correlations with NU. Similarly, in the female cohort, the QA values in the CC were positively correlated with the severity of PE and PM. Our observations regarding the correlation between the QA values in the forceps minor and major and the UPPS subscales were also divergent between males and females. This underscores the existence of sex-based disparities in the alterations of the CC among individuals with impulsivity, a phenomenon previously discussed by Silveri et al. (2006). Collectively, our findings highlight a substantial association between the alterations in the CC tract and impulsive behaviors. Nonetheless, the precise direction of these alterations remains ambiguous, necessitating further research employing more precise modalities such as DMRI to elucidate the specific white matter modifications occurring within this tract.

Another important finding of our investigations was the positive correlation between the QA values in the bilateral cingulum and PM in the entire cohort, and in the males and the females, as well as the severity of NU in the male cohort. The cingulum bundle, located above the corpus callosum, links the frontal, parietal, and temporal lobes (Wu et al., 2016) and plays a critical role in facilitating the cognitive control and the executive function governed by the frontal cortex (Bubb, Metzler-Baddeley and Aggleton, 2018). While prior reports predominantly indicate an inverse relationship between the white matter integrity in the cingulum tract and impulsivity in patients (Chiang et al., 2015; Damatac et al., 2022; Scholz et al., 2016), our study demonstrates a positive correlation in healthy individuals. Nevertheless, one of our intriguing observations was a negative correlation between the QA values in the left cingulum and the severity of NU in the female cohort. A plausible explanation for this discrepancy is the mediating function of the cingulum tract on the activities of the frontal lobe. According to this conjecture, decreased cingulum integrity could potentially lead to diminished control over the frontal cortex, thereby elevating the risk of impulsive behaviors. However, we primarily observed a positive correlation between cingulum integrity and impulsivity in healthy individuals, possibly stemming from a compensatory mechanism aimed at repressing escalated frontal cortex function. Noteworthy, our study exclusively encompassed healthy individuals, in contrast to the cited investigations which focused on psychiatric patients, particularly attention deficit hyperactivity disorder. This emphasizes the necessity of further research aimed at elucidating the intricate role of the cingulum tract in the manifestation of impulsive behaviors.

Furthermore, our subgroup analysis has unveiled additional white matter tracts whose integrity seems to be correlated with impulsivity in healthy individuals. For instance, we observed a positive correlation between the QA values in the right medial lemniscus and the severity of PE in the female cohort. This discovery is consistent with the findings of Alfano et al. (2021), who observed that healthy individuals exhibiting impulsivity tend to exhibit elevated FA values in the medial lemniscus tract (Alfano et al., 2021). The medial lemniscus is a pivotal conduit for conveying sensory spinothalamic information to the primary somatosensory cortex (Navarro-Orozco and Bollu, 2023). Thus, the positive correlation between impulsivity and the white matter integrity within this tract can be justified by the hypothesis that the heightened integrity may facilitate an augmented flow of somatosensory information to the somatosensory cortex, potentially resulting in exaggerated responses to external environmental stimuli. Analogous to the medial lemniscus tract, the ATR constitutes a white matter pathway responsible for transmitting sensory and motor information to the cerebral cortex (George and J, 2023). On the other hand, we observed a negative correlation between the QA values in the right ATR and the severity of NU in the female cohort. Based on this observation, it might be hypothesized that the ATR conveys a different set of information to the cerebral cortex, predominantly engendering inhibition of impulsive behaviors rather than their induction.

Moreover, we found a positive correlation between the QA values in the right CST and the PE in the female cohort. This was in line with previous reports of the increased integrity within the CST of healthy individuals showing an increased tendency for impulsive behaviors (Alfano et al., 2021). CST is a white matter tract that controls primary motor functions and is involved in voluntary movements (Van Wittenberghe and Peterson, 2023). Consequently, individuals with heightened CST integrity and function may exhibit a heightened susceptibility to motor impulsive behaviors. However, a deeper exploration is imperative to ascertain the precise role of the CST in impulsive behavior.

Concomitantly, an inverse relationship emerged between the white matter integrity of the fornix and the severity of sensation seeking in the male cohort. The fornix, an integral component of the limbic system situated beneath the corpus callosum, constitutes a C-shaped white matter tract along the medial aspects of the cerebral hemispheres, facilitating connectivity between the medial temporal lobes and the hypothalamus (Choi, Lee and Lee, 2021). Crucially, the fornix functions as the primary afferent and efferent pathway of the hippocampus, exerting a pivotal influence on cognitive processes (Choi, Lee and Lee, 2021). Given its role, the presence of altered fornix integrity in individuals with impulsivity is not unexpected; however, sufficient evidence on this matter is lacking. For instance, Onnink et al. (2015) observed distinct diffusivity patterns in the fornix of ADHD patients compared to their healthy counterparts (Onnink et al., 2015). Nevertheless, there is a paucity of research focusing explicitly on white matter alterations in individuals exhibiting impulsive traits and this underscores the necessity for further investigations to untangle the precise involvement of the fornix in impulsivity.

Our study possesses several notable strengths. Notably, it employs an innovative imaging analysis technique, namely DMRI connectometry, to elucidate the intricate associations between white matter microstructure and impulsivity. Furthermore, the study undertakes a comprehensive examination of the potential sex-based disparities inherent in these associations.

On the other hand, our study has several limitations. Foremost among these is the relatively modest sample size, which may curtail the generalizability of our findings. Additionally, the cross-sectional nature of the study design necessitates prudence in drawing causal inferences. While our investigation offers valuable insights, further research endeavors employing longitudinal approaches could potentially provide a more robust understanding of these dynamics.

## 5. Conclusion

There are notable correlations between impulsivity and alterations in the white matter integrity across diverse tracts, encompassing the corpus callosum, the cerebellar pathways, and the cingulum. In addition, the patterns of these correlations differ between males and females. While our study contributes valuable insights by unveiling considerable associations between microstructural modifications in white matter and impulsive behaviors of healthy individuals, further research is imperative to better understand these dynamics. By unraveling the intricacies of these neural correlates, we can pave the way for a more comprehensive knowledge of the neural basis of impulsive behavior.

## Author Contribution

**Fatemeh Rashidi:** Writing – original draft. **Mohammadamin Parsaei:** Writing – original draft, review & editing. **Iman Kiani:** Writing – original draft. **Arash Sadri:** Supervision, review & editing. **Mohammad Hadi Aarabi:** Supervision, review & editing. **Seyed Reza Darijani:** Review & editing. **Yung Sang Lee:** Supervision, review & editing. **Hossein Sanjari Moghaddam:** Conceptualization, statistical analysis, supervision, review & editing.

## Declaration of generative AI and AI-assisted technologies in the writing process

During the preparation of this work, we cautiously used AI (ChatGPT 3.5/OpenAI) only to improve the readability and language of our manuscript. After using ChatGPT 3.5/OpenAI, the authors reviewed and edited the content as needed and take(s) full responsibility for the content of the publication.

